# Collagen Nanofiber Reinforced Alginate Hydrogel Tube Microbioreactors for Cell Culture

**DOI:** 10.1101/2025.04.23.650245

**Authors:** Xinran Wu, Yakun Yang, Ying Pan, Yong Wang, Xiaojun Lian, Cheng Dong, Shue Wang, Aijun Wang, Yuguo Lei

## Abstract

The large-scale production of mammalian cells is pivotal for various applications; however, current bioreactor technologies encounter significant technical and economic challenges. Scaling up cell cultures remains problematic due to excessive cell aggregation, shear stress-induced cell death, batch-to-batch inconsistencies, and limited scalability. We propose that engineering a cell-friendly microenvironment can enhance culture efficiency. Previously, we developed alginate hydrogel microtubes (AlgTubes) that significantly improved cell density and growth rates. Nevertheless, AlgTubes lack adhesion sites essential for anchorage-dependent cells and frequently break, causing cell leakage and production inconsistencies. To address these limitations, this study reinforced AlgTubes with collagen nanofibers, creating collagen-alginate hybrid hydrogel microtubes (ColAlgTubes). Utilizing a novel micro-extruder, we efficiently produced cell-loaded ColAlgTubes. Collagen formed a dense nanofiber network interwoven with the alginate mesh, enhancing the hydrogel’s mechanical properties while providing cell adhesion sites. ColAlgTubes protected cells from hydrodynamic stress and maintained cell mass within a 400 μm diameter, ensuring efficient nutrient exchange and waste removal. This optimized microenvironment resulted in high cell viability, rapid proliferation, and exceptional yields of 5×10^8^ cells/mL - 200 times higher than conventional culture methods. With their scalability, cost-effectiveness, and efficiency, ColAlgTubes offers a transformative solution for large-scale cell production with broad applications in biotechnology, regenerative medicine, and therapeutic manufacturing.

## A. Introduction

Human and animal cells are integral to a broad spectrum of biomedical and industrial applications. Stem cells, such as human pluripotent stem cells (hPSCs) including human embryonic stem cells (hESCs) and induced pluripotent stem cells (iPSCs), along with their differentiated progeny, are utilized in the treatment of various diseases and injuries^1^. They are also essential for disease modeling, drug screening, and toxicity testing^1^. Immune cells, like T cells and natural killer cells, are extensively used in cancer immunotherapy^2–6^. Additionally, mammalian cells are crucial for producing recombinant proteins and viruses, which are vital for research and clinical applications^7–9^.

These applications need large quantities of high-quality cells^1^. For example, treating a single patient requires approximately 10^9^ cardiomyocytes for myocardial infarction (MI) or 10^9^ β cells for Type 1 diabetes^10^. The demand is further amplified by the prevalence of degenerative diseases and organ failure, with 1–2.5 million people in the U.S. affected by Type 1 diabetes and around 8 million by MI^10^. Similarly, tissue engineering demands large numbers of cells, with approximately 10^10^ hepatocytes or cardiomyocytes needed to construct an artificial human liver or heart^11^. Drug discovery efforts, such as screening a million-compound library, can also require 10^10^ cells per screen^10^. Moreover, large quantities of mammalian cells, such as Chinese Hamster Ovary (CHO) cells and Human Embryonic Kidney 293 (HEK293) cells, are essential for producing therapeutic biologics, including monoclonal antibodies, enzymes, and viral particles^7–9^.

Despite the critical need for efficient and scalable cell production, existing methods remain inadequate, particularly for clinical applications^1,10,12^. Two-dimensional (2D) culture systems, such as flasks, are widely used but lack the complexity of natural cellular environments. They are also resource-intensive, requiring significant labor, space, and reagents, making them impractical for large-scale production. Three-dimensional (3D) suspension culture systems, such as stirred-tank bioreactors, have been developed to improve scalability^13,14^. However, these systems face significant challenges, particularly uncontrolled cell aggregation. Aggregates exceeding 400 μm in diameter suffer from impaired transport of nutrients, oxygen, and growth factors, leading to slower proliferation, apoptosis, and undesired differentiation. While agitation can mitigate aggregation, it also introduces shear forces, negatively impacting cell survival, growth, and differentiation efficiency^10,15,16^. As a result, 3D suspension cultures often exhibit high cell mortality, slow proliferation rates, and low volumetric yields. For instance, hPSCs in stirred-tank bioreactors typically undergo only a four-fold expansion over four days, yielding approximately 2.0×10^6^ cells/mL, which utilizes just 0.4% of the bioreactor volume^17–19^.

The complexity of hydrodynamic conditions in stirred-tank bioreactors - such as flow dynamics, shear stress, and chemical gradients - adds further challenges. These factors depend on variables including bioreactor design, medium viscosity, and agitation speed, making precise control difficult^1,10,12,15–20^. The variable hydrodynamic conditions contribute to large cell production variation. For example, in cardiomyocyte production from hPSCs, yields from three ∼100 mL batches of hESCs varied widely, ranging from 40 to 100 million cells, with cardiomyocyte purity fluctuating between 54% and 84%^21,22^. When using a different hPSC line under the same conditions, yields ranged from 89 to 125 million cells, with purity varying from 28% to 88%^21,22^. Scaling production from 100 mL to 1000 mL requires extensive re-optimization of agitation rates and culture protocols, underscoring the technical and economic challenges of achieving industrial-scale cell manufacturing^21,22^. Currently, the largest demonstrated hPSC suspension cultures are limited to volumes of just tens of liters^10,23^.

To overcome these limitations, we propose the development of hydrogel tube-based 3D microbioreactors. Our previous studies demonstrated that culturing cells within hollow hydrogel microtubes or microbioreactors made from alginate polymers mitigate many of these challenges. These microtubes prevent excessive cell aggregation, enhance mass transport efficiency, and eliminate hydrodynamic stress, leading to high cell viability, rapid proliferation, and significantly improved volumetric yields. Notably, yields of up to 5×10^8^ cells/mL have been achieved^24–33^. These findings highlight the transformative potential of hydrogel tube microbioreactors for scalable, efficient, and cost-effective cell production^24–33^.

Despite their advantages, alginate hydrogel microtubes have critical limitations. They are mechanically fragile, with a significant proportion breaking during cell culture, leading to cell leakage and production failures. Additionally, many cells do not grow well in alginate hydrogel microtubes because they cannot adhere to the alginate surface. There is a critical need for new, robust, and adhesive hydrogel tube microbioreactors. This study introduces the fabrication and application of hybrid collagen-alginate microtubes for cell culture. Within these microtubes, collagen proteins form a nanofiber network that interpenetrates the alginate hydrogel mesh. The collagen nanofibers enhance the alginate hydrogel’s structural integrity and serve as anchoring points for adhesion-dependent cell growth. This advanced system will facilitate cost-effective large-scale cell culture for various biomedical and industrial applications.

## B. Methods

### Collagen Extraction from Rat Tails

Rat tails were harvested and soaked in 70% ethanol for 30 minutes. The skin was carefully removed using a scalpel and forceps to isolate the tendons, which were then rinsed three times with PBS. To ensure sterilization, the tendons were submerged in 70% ethanol for at least one hour. After sterilization, they were soaked in 0.02N acetic acid and stirred at 4°C for 48 hours. The resulting viscous mixture was centrifuged at 10,000 rpm for 60 minutes at 4°C to remove debris. The supernatant was collected and dialyzed against 0.02N acetic acid, yielding the collagen stock solution for further applications.

### Process ColAlgTubes

Rat tail collagen was adjusted to pH 5.0 using a neutralizing solution and then combined with alginate in a pH 5.0 NaCl solution to create a collagen-alginate mixture. A custom-built micro-extruder was used to fabricate ColAlgTubes. A 2% methylcellulose (MC) solution containing single cells was pumped into the central channel at a rate of 20 μL/min, while a pre-cooled syringe containing the collagen-alginate solution was placed in a homemade ice box and pumped into the side channel at 61 μL/min. The two solutions were co-extruded into a 50 mM HEPES + 100 mM CaCl_2_ buffer (pH 7.4), forming ColAlgTubes. After extrusion, the buffer was replaced with a cell culture medium to support cell growth.

### Scanning Electron Microscope (SEM) Imaging

The nanostructures of ColAlgTubes were characterized using a Zeiss SIGMA VP-FESEM. Before imaging, samples were dehydrated through a graded ethanol series (25%, 50%, 70%, 85%, 95%, and 100%), with each step lasting 5 minutes. Critical point drying was performed using a Leica EM CPD300 Critical Point Dryer. The samples were sputter-coated with a 4.5 nm iridium layer to enhance conductivity using a Leica EM ACE600 Sputter Coater. SEM images were acquired at an accelerating voltage of 5 kV, with a working distance of 6 mm and magnifications ranging from 150x to 5,000x, allowing detailed visualization of the nanofiber structures.

### Collagen and Alginate Degradation

ColAlgTubes were incubated with 0.2 mg/mL Collagenase P at 37°C for 15 minutes to degrade collagen nanofibers. ColAlgTubes were incubated with 0.5 mM EDTA at room temperature for 15 minutes to dissolve the alginate hydrogel. Following degradation, the samples were prepared for SEM imaging as required.

### Culturing hPSCs in Collagen Alginate Tubes

For a typical cell culture, 20 μL of cell solution in ColAlgTubes was suspended in 2 mL of E8 medium supplemented with 10 μM Y-27632 in a 6-well plate. The culture was maintained in an incubator at 37 °C with 5% CO_2_ and 21% O_2_, and the medium was refreshed daily. For cell passaging, the medium was removed, and ColAlgTubes were dissolved using 0.2 mg/mL Collagenase P containing 0.5 mM EDTA for 20 minutes. The cell mass was collected by centrifugation at 100 g for 5 minutes, followed by treatment with Accutase at 37 °C for 10 minutes. The cells were then dissociated into single cells for subsequent culture or cryopreservation.

### Cardiomyocyte differentiation

Human pluripotent stem cells (hPSCs) in ColAlgTubes were cultured in E8 medium supplemented with 10 μM Y-27632 until the diameter of the cell aggregates reached the inner diameter of the ColAlgTubes. On Day 0, the medium was replaced with E5 medium containing 10% lipid mix and 5 μM CHIR9902. After 24 hours, the medium was replaced with E5 medium containing 10% lipid mix and 3 μg/mL Heparin. From Day 2 to Day 4, the medium was refreshed daily with E5 medium containing 10% lipid mix, 3 μg/mL Heparin, and 3 μM IWR1. From Day 5 to Day 6, the medium was refreshed daily with E5 medium containing 10% lipid mix and 3 μg/mL Heparin. From Day 7 to Day 10, the medium was refreshed every other day with an E5 medium containing 10% lipid mix and 20 μg/mL insulin. From Day 11 to Day 18, the medium was refreshed every other day with DMEM medium (without glucose, glutamine, and pyruvate) supplemented with 5 mM Lactate. Cardiomyocytes were collected on Day 18 for analysis. For long-term culture, the medium was switched to α-MEM supplemented with 5% FBS and refreshed every other day.

### Staining, Flow Cytometry, and Imaging

Single-cell suspensions were fixed in 4% paraformaldehyde (PFA) at room temperature for 15 minutes. The fixed cells were then incubated overnight at 4°C with primary antibodies in PBS containing 0.1% Triton X-100 and 0.5% BSA. After extensive washing, secondary antibodies were added and incubated for 2 hours at room temperature. The cells were washed three times with PBS supplemented with 0.5% BSA before analysis using the Attune® NxT™ Acoustic Focusing Cytometer (ThermoFisher). Flow cytometry data were processed using FlowJo software. LIVE/DEAD® Cell Viability staining was performed following the manufacturer’s instructions for cell viability assessment. Phase-contrast and fluorescence imaging were conducted using a Zeiss Axio Observer Fluorescent Microscope.

### Statistical Analysis

Data analysis was conducted using GraphPad Prism 8, with results presented as mean ± standard error of the mean (SEM). Statistical significance was determined using appropriate tests based on the type of comparison: One-way analysis of variance (ANOVA) was used for multiple group comparisons (three or more groups). Unpaired two-tailed t-tests were performed for comparisons between the two groups. Statistical significance was indicated as follows: p < 0.05 (*), p < 0.01 (**), p < 0.001 (***).

## C. Results

### Fabrication of Collagen-Alginate Hybrid Hydrogel Tubes (ColAlgTubes)

This study aimed to develop ColAlgTubes as an innovative cell culture platform. Currently, no technology exists for rapidly fabricating hybrid ColAlgTubes for cell culture applications. To address this gap, we designed a specialized micro-extruder with dual inlets and a single outlet (Fig. 1A). The extruder was constructed using a Formula 3B printer and clear resin. Additionally, we designed and 3D-printed a cooling box to maintain the collagen-alginate solution at a low temperature (Fig. 1B). The ColAlgTube extrusion setup comprises two syringe pumps, the micro-extruder, the cooling box, and a container filled with HEPES buffer (Fig. 1C, D). Syringe 1 is loaded with a room-temperature (RT) cell solution, while Syringe 2 contains an ice-cold collagen-alginate mixture (pH = 5.0). The cooling box features a designated channel to hold Syringe 2 and is packed with ice, ensuring the collagen-alginate solution remains below 4°C (Fig. 1B, C). Maintaining a low temperature and acidic pH prevents premature collagen gelation inside the syringe.

**Figure 1.**
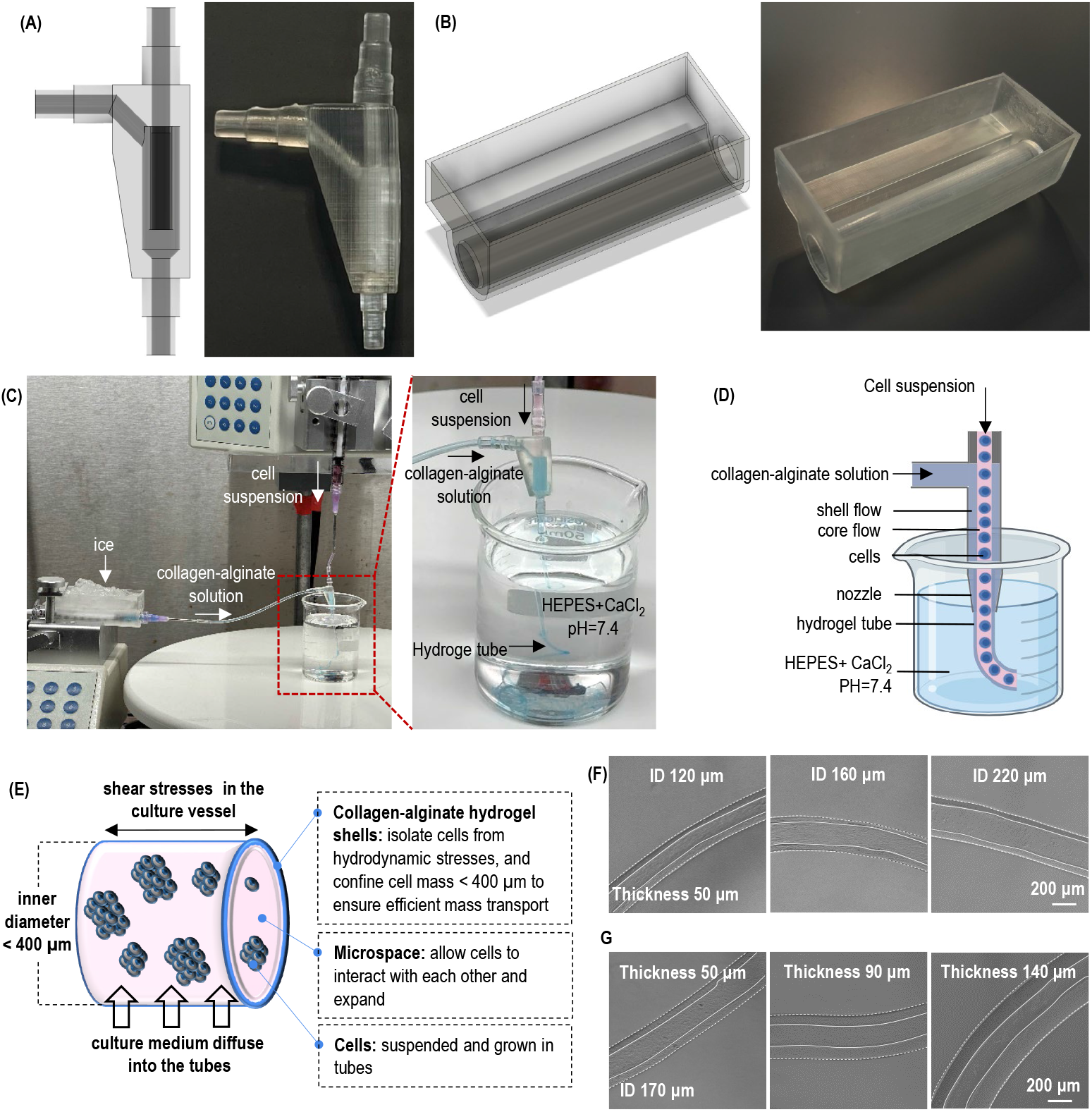
The Collagen-Alginate Hydrogel Tube (ColAlgTube) Microbioreactors. **(A)** Design and printed version of the new micro-extruder. **(B)** Design and printed version of the cooling box used for temperature control. **(C, D)** The setup for processing ColAlgTubes includes two syringe pumps, a custom-made micro-extruder, a cooling box, and a HEPES buffer reservoir. To produce ColAlgTubes, a cell solution and an ice-cold collagen-alginate solution (pH = 5.0) are pumped into the central and side channels of the micro-extruder, respectively, forming coaxial core-shell flows. These flows are extruded into a HEPES-CaCl_2_ buffer (pH = 7.4), which neutralizes the collagen-alginate solution. The shell flow forms a hydrogel tube through collagen fiber formation at pH = 7.4 and alginate crosslinking via Ca^2+^ ions. **(E)** Cells are cultured within ColAlgTubes suspended in the culture medium inside a vessel. The tubes protect cells from hydrodynamic stress and confine the cell mass to a radial diameter of less than 400 μm, ensuring efficient mass transport of nutrients and waste. These hydrogel tubes create a cell-friendly microenvironment, allowing efficient cell interaction, growth, and diffusion of nutrients through the hydrogel shell. **(F)** The inner diameter (ID) is varied from 120 to 220 μm while maintaining a shell thickness of 50 μm. **(G)** The shell thickness is adjusted from 50 to 140 μm while keeping the ID constant at 170 μm.

To fabricate ColAlgTubes, the two solutions are pumped into inlets 1 and 2, generating coaxial core-shell flows that are extruded into a HEPES buffer solution containing 100 mM CaCl_2_ at 37°C and pH 7.4 (Fig. 1C, D). The core flow carries the cell solution, while the shell flow contains the collagen-alginate mixture. At the microscale, these solutions establish stable laminar flows. The acidic collagen solution in the shell flow is neutralized by the core cell solution and the HEPES buffer, triggering the rapid formation of a collagen nanofiber network. Simultaneously, the alginate in the shell undergoes crosslinking with Ca^2+^ ions, forming a stable alginate network. This is the first time to rapidly process an interpenetrating collagen and alginate tube for cell culture.

### Engineering Principles and Control ColAlgTube Dimensions

Similar to our previously developed AlgTubes, ColAlgTubes overcome the limitations of conventional cell culture techniques by providing a physiologically relevant microenvironment for cell growth (Fig. 1E). Cells are cultured within microscale ColAlgTubes, which remain suspended in the culture medium. The hydrogel tubes create free spaces for cell-cell interactions and expansion while protecting cells from hydrodynamic stresses within the culture vessel. Additionally, the tubes restrict cell mass to a radial diameter of less than 400 μm, aligning with human tissue’s diffusion limit, ensuring efficient nutrient exchange and waste removal throughout the culture. Moreover, collagen nanofibers within the ColAlgTubes serve as adhesive substrates, supporting the growth of anchor-dependent cells.

The dimensions of ColAlgTubes can be tuned by adjusting the ratio of core and shell flows, as well as the diameter of the extruder nozzle. Wall thickness can be controlled by modifying the collagen-alginate shell flow rate, with lower flow rates producing thinner walls. Figures 1F and G showcase ColAlgTubes with varying diameters and wall thicknesses, highlighting the adaptability of this system.

### Nanostructures of ColAlgTubes

We hypothesize that (1) collagen and alginate form an interpenetrating network and (2) collagen nanofibers reinforce the alginate hydrogel, enhancing the mechanical strength of ColAlgTubes and increasing their resistance to force-induced breakage. We used scanning electron microscopy (SEM) to examine the nanostructures of ColAlgTubes. ColAlgTubes were fabricated with iPSC cells and cultured until the tubes were partially filled. The tubes were then fixed with paraformaldehyde (PFA), dehydrated through a graded ethanol series, metal-sputtered, and imaged using SEM. We captured images of the overall tube structure, inner and outer surfaces, wall nanostructures, and encapsulated cells. The alginate concentration was maintained at 0.75%, while collagen concentrations varied from 1 to 3 mg/mL (Fig. 2A–C). For comparison, pure alginate and pure collagen hydrogel tubes were also prepared (Fig. 2D, E).

**Figure 2.**
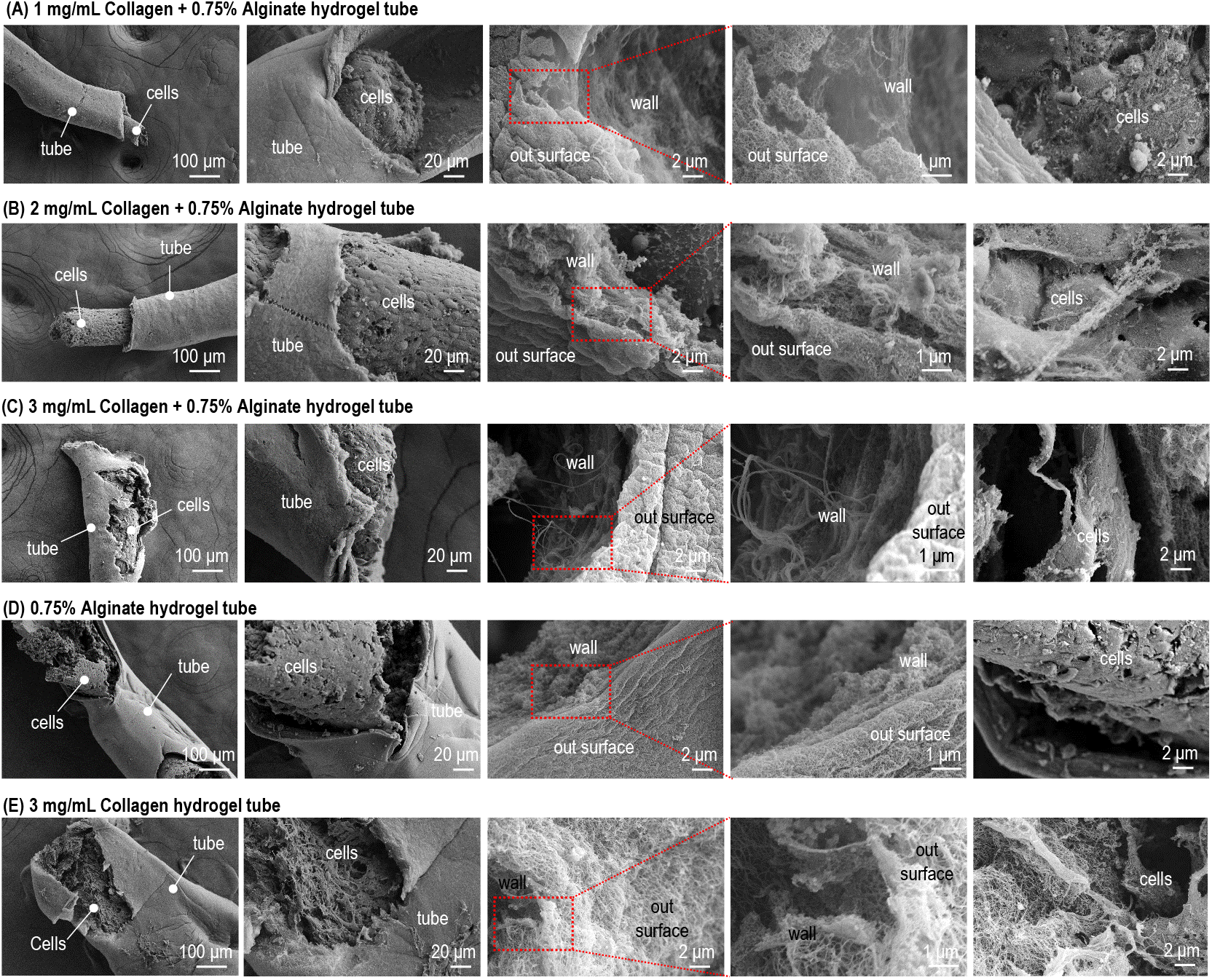
Nanostructures of ColAlgTubes Containing iPSCs. **(A)** 1 mg/mL Collagen + 0.75% Alginate hydrogel tube. **(B)** 2 mg/mL Collagen + 0.75% Alginate hydrogel tube. **(C)** 3 mg/mL Collagen + 0.75% Alginate hydrogel tube. **(D)** 0.75% Alginate-only tube. **(E)** 3 mg/mL Collagen-only tube.

In pure alginate hydrogel tubes, a dense mesh-like structure was observed (Fig. 2D), while pure collagen hydrogel tubes were composed of collagen nanofibers (Fig. 2E). The hybrid ColAlgTubes exhibited nanostructures similar to alginate hydrogel tubes but contained small numbers of collagen nanofibers interwoven within the dense alginate matrix (Fig. 2A–C). Increasing the collagen concentration from 1 mg/mL to 3 mg/mL resulted in more collagen nanofibers.

To further confirm the interpenetrating network, alginate was selectively dissolved using an EDTA solution, allowing clear visualization of the collagen nanofiber network (Fig. 3A, B). Conversely, collagen nanofibers were degraded with collagenase, revealing the dense alginate network (Fig. 3C). These findings validate our hypothesis that ColAlgTubes consist of an interpenetrating network of collagen nanofibers and alginate hydrogels.

**Figure 3.**
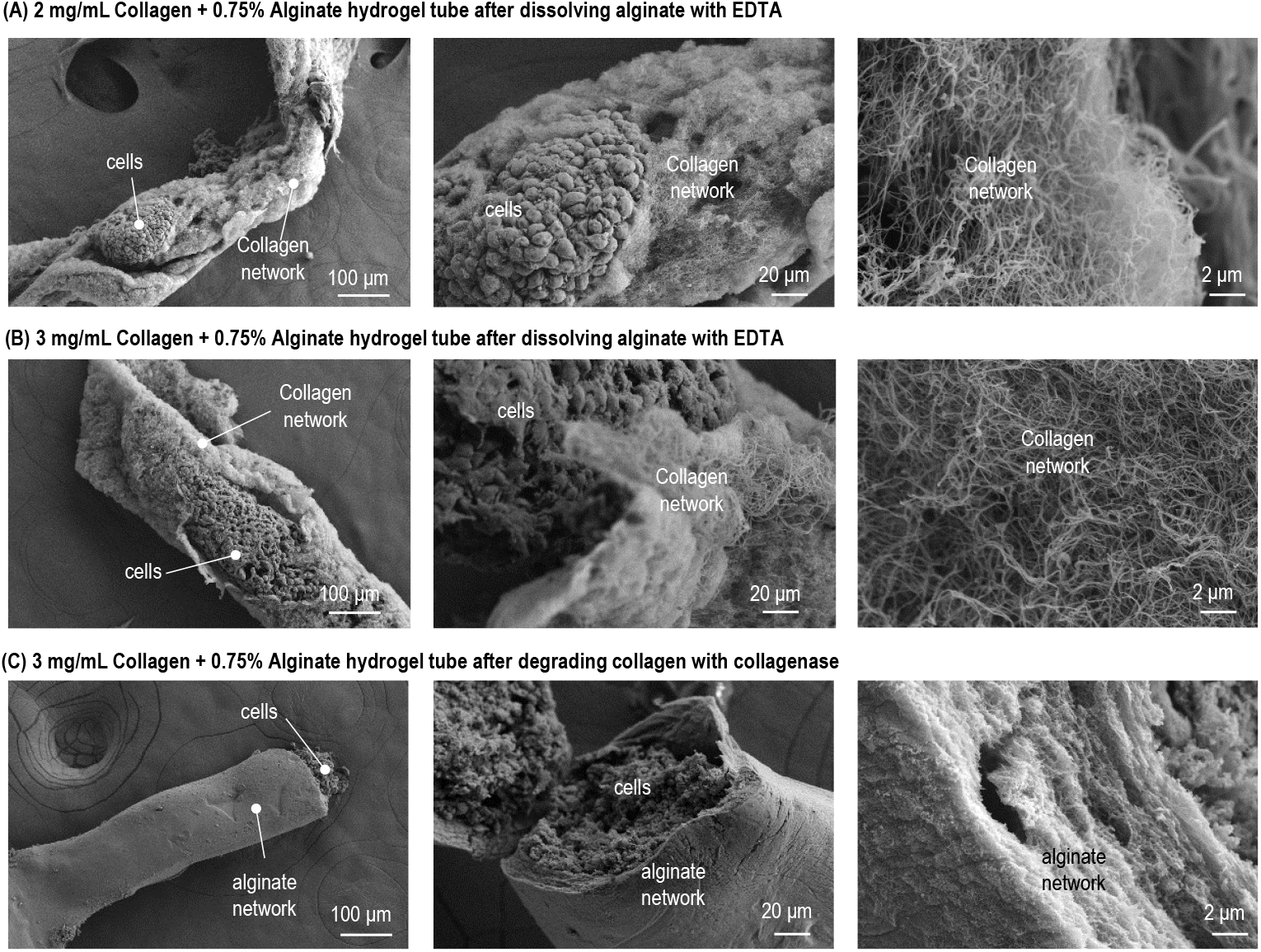
Nanostructures of ColAlgTubes Containing iPSCs. **(A)** 2 mg/mL Collagen + 0.75% Alginate hydrogel tube after dissolving alginate with EDTA. **(B)** 3 mg/mL Collagen + 0.75% Alginate hydrogel tube after dissolving alginate with EDTA. **(C)** 3 mg/mL Collagen + 0.75% Alginate hydrogel tube after degrading collagen with collagenase.

### Culturing HEK293 Cells in ColAlgTubes

To assess the suitability of ColAlgTubes for cell culture, HEK293 cells (Fig. 4A) were processed into the tubes. Within 24 hours, the cells adhered to the inner surface and began expanding into colonies. Over time, they proliferated further, eventually filling the entire tube and forming a three-dimensional (3D) cell mass. Live/Dead cell staining confirmed that most cells remained viable. Over 4 × 10^8^ cells/mL were successfully harvested. In contrast, when HEK293 cells were cultured in pure AlgTubes, they did not adhere to the tube walls. Instead, they formed unattached spheroids (Fig. 4B). Their growth rate was significantly lower compared to those cultured in ColAlgTubes, highlighting the importance of adhesion for HEK293 cell proliferation (Fig. 4B). These results confirm our hypothesis that incorporating collagen fibers into AlgTubes provides essential anchoring points for cell attachment and growth.

**Figure 4.**
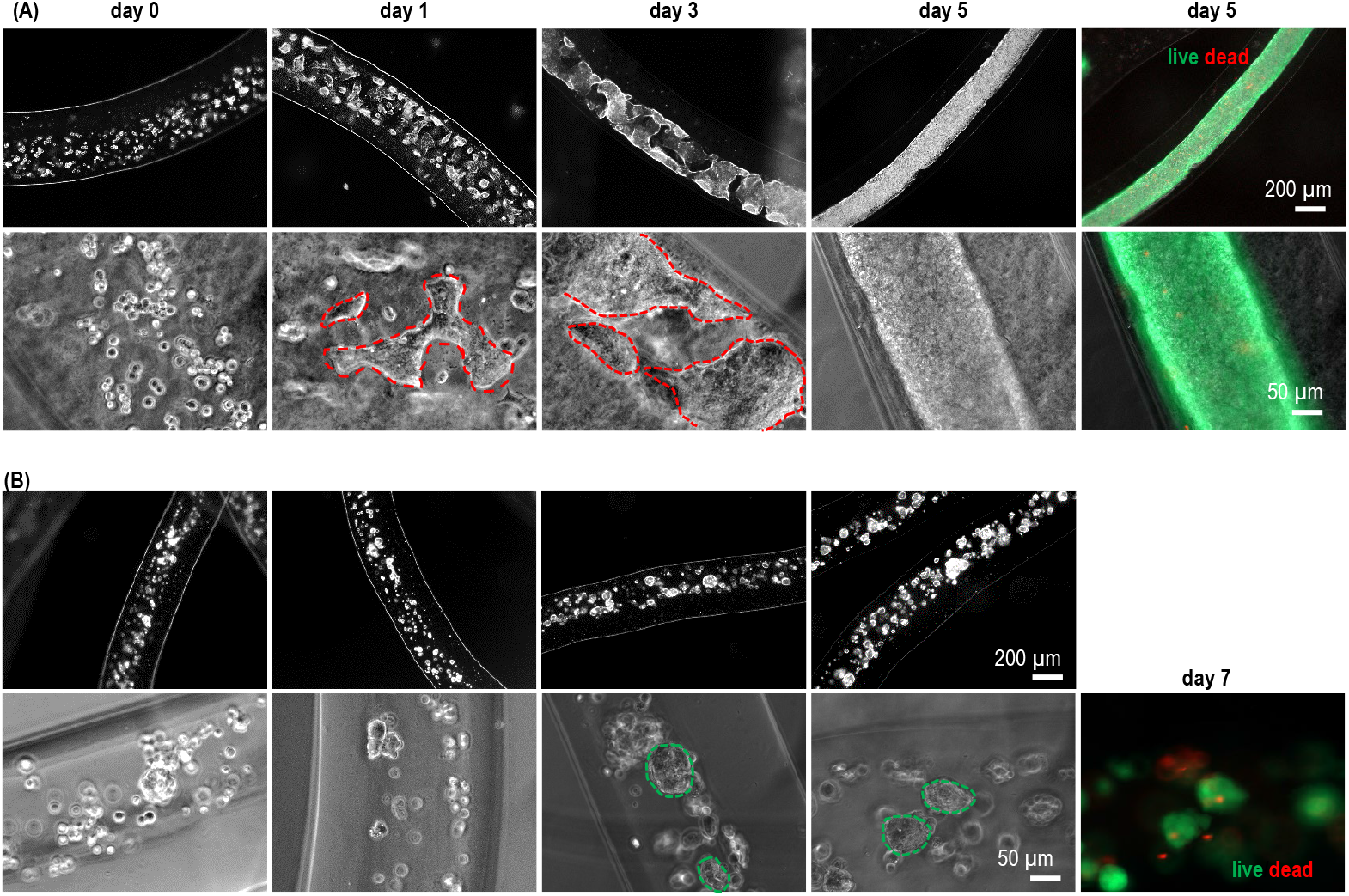
Culturing HEK 293T Cells in ColAlgTubes and AlgTubes. Phase-contrast and Live/Dead staining images of 293T cells cultured in ColAlgTubes **(A)** and AlgTubes **(B)** at various time points. In ColAlgTubes, 293 cells adhered to inner surface of the tubes as outlined with the red dash lines in (A), while in AlgTubes, 293 cells did not attach to the tube and grew as unattached spheroids as outlined with the green dash lines in (B).

### Culturing Human Pluripotent Stem Cells (hPSCs) in ColAlgTubes

H9 human embryonic stem cells (hESCs) were cultured within ColAlgTubes. Within 24 hours, the cells adhered to the inner surface of the tubes. By day 3, small colonies had formed, which gradually developed into spheroids by day 5. By day 9, cell proliferation had expanded to fill most of the tube (Fig. 5A). Live/Dead staining performed before and after cell release indicated minimal cell death (Fig. 5C, D). The dissociated cell mass was subsequently fixed with paraformaldehyde (PFA) and stained for the pluripotency markers Nanog and Oct4. Flow cytometry analysis confirmed that 97.0% and 98.3% of the cells expressed Nanog and Oct4, respectively, demonstrating that hPSCs retained their pluripotency after culturing in ColAlgTubes (Fig. 5E). Additionally, more than 4.5 × 10^8^ cells per milliliter of microspaces were harvested. In contrast, when hPSCs were cultured in pure AlgTubes (Fig. 5B), they did not adhere to the tube and grew as unattached spheroids. This further supports the role of collagen fibers in providing anchoring points essential for cell attachment and growth.

**Figure 5.**
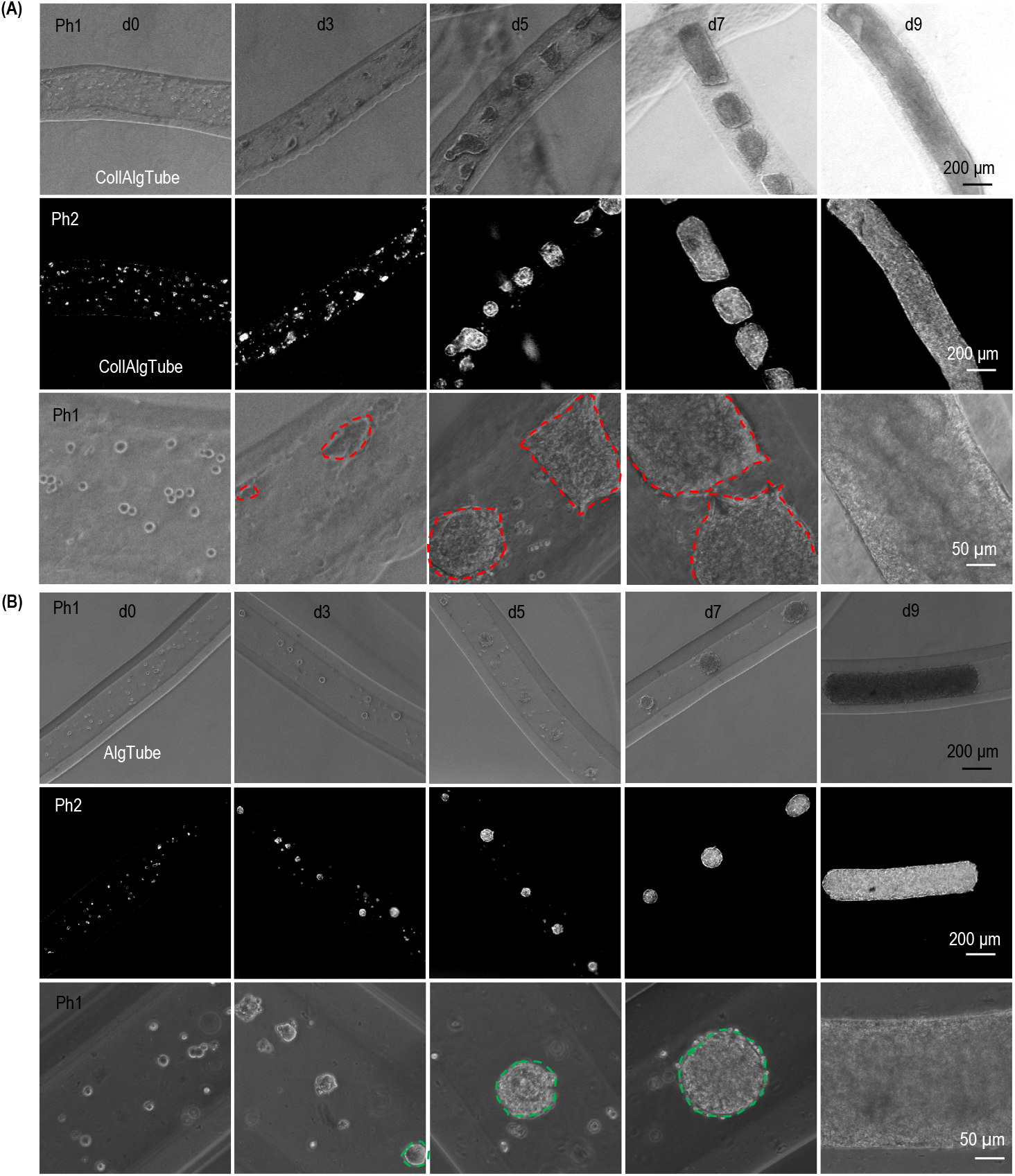
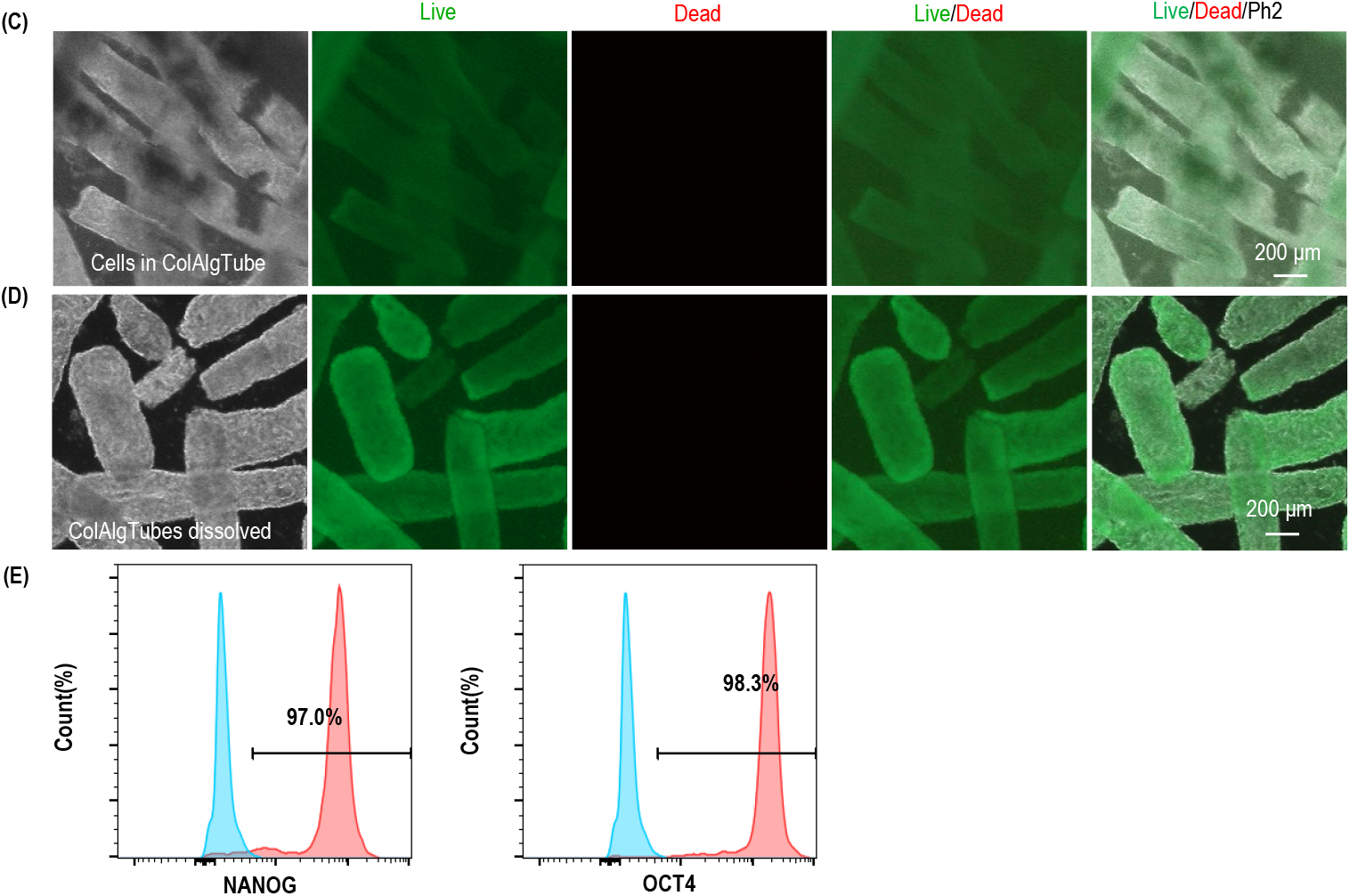
Culturing H9 hESCs in ColAlgTubes. **(A)** Phase-contrast (Ph1) and darkfield (Ph2) images of H9 hESCs cultured in ColAlgTubes on days 0, 1, 3, 5, 7, and 9. Cells adhered to inner surface of ColAlgTubes as shown by the red dash lines. **(B)** Phase-contrast (Ph1) darkfield (Ph2) images of human iPSCs cultured in AlgTubes on days 0, 1, 3, 5, 8 and 9. did not attach to the tube and grew as unattached spheroids as shown by the green dash lines. **(C, D)** Live/Dead staining of day nine H9 cells within ColAlgTubes (C) and after release from ColAlgTubes (C). **(E)** Flow cytometry analysis showing that the majority of cells harvested on day 9 express pluripotency markers OCT4 and Nanog.

### Differentiating hPSCs into Cardiomyocytes in ColAlgTubes

To further explore the potential of ColAlgTubes, we differentiated hPSCs into cardiomyocytes. H9 hESCs were cultured in ColAlgTubes and expanded in Essential 8 (E8) medium for seven days. Without requiring passaging, the medium was switched to a mesoderm induction medium to initiate differentiation into mesoderm progenitors. On day 2, the medium was replaced with cardiac progenitor differentiation medium, followed by cardiomyocyte differentiation medium on day 7. From days 11 to 18, cells were maintained in a metabolic enrichment medium, with the majority remaining viable. Following treatment with EDTA and collagenase, fibrous cardiac tissues were successfully harvested (Fig. 6A). Further dissociation of these tissues using Accutase yielded over 3 × 10^8^ cardiomyocytes per milliliter of hydrogel tubes. “Immunostaining revealed that most cells expressed the cardiomyocyte marker cTnT, demonstrating that ColAlgTubes effectively support stem cell differentiation.

**Figure 6.**
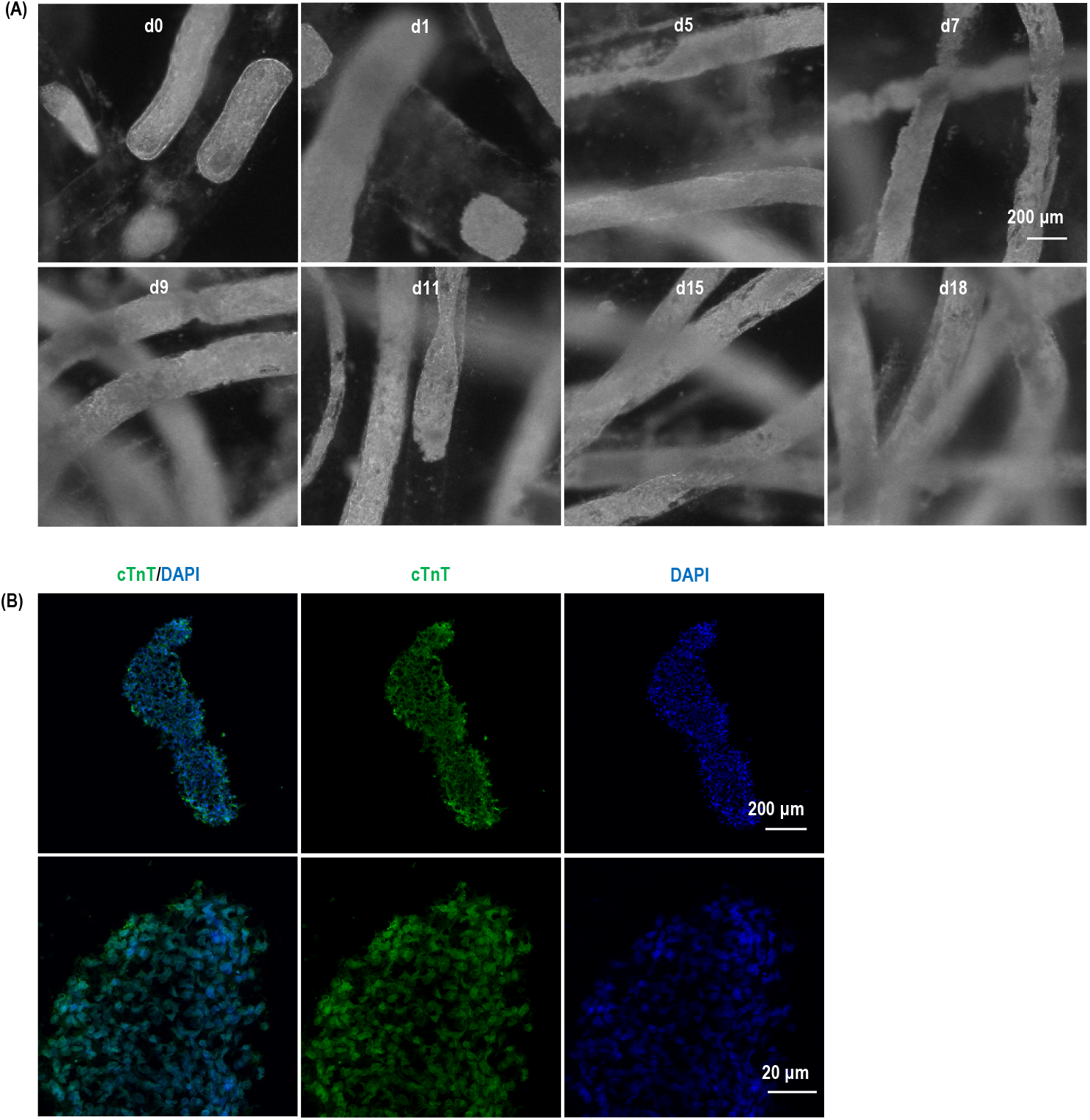
Differentiation of H9 hESCs into Cardiomyocytes in ColAlgTubes. **(A)** Phase-contrast of H9 derived cardiomyocytes cultured in ColAlgTubes at days 0, 1, 5, 7, 9, 11, 15, and 18. **(B)** Immunostaining of cTnT marker for cardiomyocytes differentiated in ColAlgTubes.

### Reduced Cell Leakage in ColAlgTubes Compared to AlgTubes

Previously, we utilized alginate-based hydrogel tubes (AlgTubes) for cell culture; however, frequent tube breakages were observed (Fig. 7A). Since cells did not adhere to AlgTubes, a significant number of cells leaked into the medium through these breaks on days 5, 6, and 7, leading to the formation of large cell aggregates (Fig. 7B). In contrast, ColAlgTubes demonstrated significantly greater structural integrity, exhibiting fewer breakages than AlgTubes. Additionally, due to their ability to support cell adhesion, cell aggregates remained attached to the tubes and did not leak into the medium (Fig. 7A, C). Quantitative analysis of leakage events (Fig. 7D) and the percentage of leaked cells on day 7 (Fig. 7E) further confirmed that ColAlgTubes had substantially lower cell leakage than AlgTubes.

**Figure 7.**
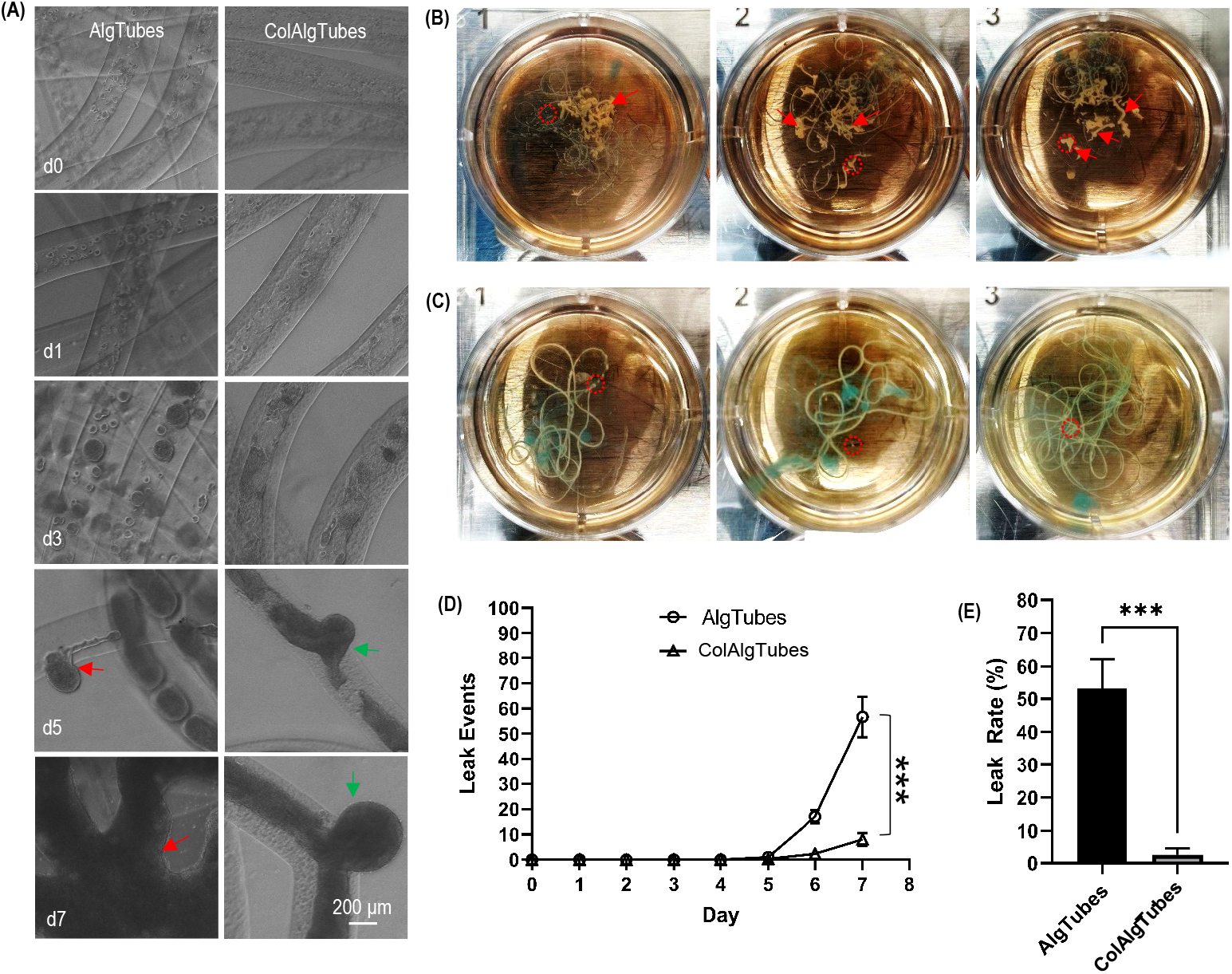
ColAlgTubes Exhibit Fewer Cell Leakage Events Compared to AlgTubes. **(A)** Phase-contrast images of H9 hESCs expanded in AlgTubes and ColAlgTubes for 7 days. Red and green arrows show cell leakage events in AlgTubes and ColAlgTubes, respectively. **(B, C)** Images showing significant cell leakage from AlgTubes, forming large cell aggregates (red arrow) (B), whereas minimal cell leakage is observed from ColAlgTubes (C). **(D)** Quantification of cell leakage events over time for AlgTubes and ColAlgTubes. **(E)** Comparison of leak rates between AlgTubes and ColAlgTubes. Leak rate is defined as the number of cells leaked into the medium divided by the total number of cells in the well. ***: p<0.001

In summary, we have demonstrated that ColAlgTubes can be a robust cell expansion and differentiation platform. The interpenetrating collagen-alginate network enhances mechanical stability, significantly reducing cell leakage compared to AlgTubes. Additionally, ColAlgTubes facilitate cell adhesion, making them suitable for culturing both adherent and suspension cells.

## D. Discussion

Current 2D and 3D cell culture methods face significant challenges in achieving efficient, scalable, and cost-effective large-scale cell production^34–36^. Key limitations include low cell yields, restricted scalability, high operational costs, and substantial culture variability^34–36^. A fundamental issue is that conventional systems, such as 2D flasks and 3D stirred-tank bioreactors, fail to accurately replicate the complex 3D microenvironments found in vivo^34–36^. In their native state, human cells exist within intricate 3D microenvironments that facilitate critical interactions with the extracellular matrix (ECM), support efficient nutrient and oxygen exchange, and minimize hydrodynamic stress^13,14,37–39^. We hypothesize that developing advanced culture technologies closely mimicking physiologically relevant microenvironments can overcome these challenges and significantly enhance culture efficiency^24,27^.

Hydrogel tube microbioreactors represent a promising approach to creating cell-friendly environments suitable for large-scale culture (Fig. 1E)^24,27^. We previously developed a method for fabricating hydrogel tubes using alginate polymers^24–33^. Alginate is abundant, cost-effective, biocompatible, and has an established clinical safety profile. It can be rapidly crosslinked with calcium ions, allowing scalable production of alginate hydrogel tubes (AlgTubes) via extrusion. After culture, AlgTubes can be dissolved with EDTA to harvest cells. Their transparency also enables real-time monitoring^24–33^.

Using AlgTubes, we cultured human pluripotent stem cells (hPSCs) over multiple passages with high consistency, maintaining pluripotency and chromosomal stability^24^. Cultures achieved high viability and rapid expansion (e.g., a 1000-fold increase over 10 days), with volumetric yields of ∼5×10^8^ cells/mL -over 200-fold higher than stirred-tank bioreactors. Furthermore, hPSCs differentiated efficiently into endothelial cells^25,28^, vascular smooth muscle cells^31^, neural stem cells^26^, and neurons^29^, all reaching yields of 5×10^8^ cells/mL. Adult cells, such as T cells, were also successfully cultured in AlgTubes. These findings highlight the transformative potential of hydrogel microtubes for high-efficiency, large-scale production^27^. Despite their success, AlgTubes have limitations. Mammalian cells lack receptors for alginate, preventing adhesion to tube walls, which limits the culture of anchorage-dependent cells like mesenchymal stem cells. Additionally, AlgTubes are mechanically fragile and prone to breakage, leading to cell leakage and variability (Fig. 7A, B). These limitations necessitate further advancements in hydrogel tube design for more robust and versatile microbioreactor systems.

Research has shown that incorporating nanofibers into hydrogels is a promising strategy for enhancing mechanical properties and structural integrity^40–42^. Nanofibers, derived from biopolymers such as collagen and silk fibroin or synthetic materials like polycaprolactone (PCL), can form an interpenetrating network within the hydrogel matrix. This structural reinforcement significantly improves the hydrogel’s tensile strength, elasticity, and stability. Additionally, nanofibers enhance cell adhesion, proliferation, and controlled drug release. By integrating nanofibers into hydrogel systems, researchers can develop advanced biomaterials with superior structural integrity and tunable mechanical properties, broadening their applications in regenerative medicine and biotechnology^40–42^.

Specific to alginate hydrogels, researchers have explored various nanofibers for reinforcement^43^. For instance, one study developed laminated composite scaffolds by combining alginate hydrogels with polycaprolactone (PCL) and gelatin electrospun mats^44^. These scaffolds demonstrated enhanced mechanical properties and controlled biodegradability, making them well-suited for cartilage tissue engineering. Another investigation created a three-dimensional composite scaffold of alginate hydrogels, PCL/gelatin nanofibers, and exosomes, effectively promoting regeneration in rat tympanic membrane perforations^45^. A recent review summarized various approaches for developing alginate composite hydrogels^43^.

However, incorporating nanofibers into alginate hydrogel tubes presents significant challenges compared to embedding nanofibers within bulk hydrogels. To the best of our knowledge, this has not been achieved by others. In this study, we developed an innovative approach to fabricate interpenetrating collagen-alginate hydrogel tubes. By integrating a cooling box, a two-flow micro-extruder, and a buffer exchange system (Fig. 1), we successfully established a method for processing collagen-alginate hybrid microtubes. Collagen proteins formed a dense nanofiber network reinforcing AlgTubes (Fig. 2, 3). As a result, ColAlgTubes exhibited remarkable durability, with significantly fewer breakages during cell culture, effectively preventing cell leakage (Fig. 7). Furthermore, since most cells express receptors for Collagen, they naturally adhered to the ColAlgTubes, enhancing cell attachment and growth (Fig. 4, 5).

Our data showed that the cell growth rate, viability, and volumetric yield achieved with ColAlgTubes were comparable to those observed with AlgTubes^24^. The high growth rate and cell density attainable in this system have significant implications for large-scale cell production. By significantly increasing cell culture density, ColAlgTubes can substantially reduce culture volume, lowering labor requirements, reagent costs, equipment needs, facility space, and manufacturing expenses.

In summary, ColAlgTubes represents a significant advancement in cell culture technology. Their ability to support high-yield cell growth and streamline large-scale manufacturing makes them a compelling alternative to conventional two-dimensional and three-dimensional culture systems. As advancements in stem cell differentiation protocols continue, future efforts should focus on integrating optimized differentiation processes into ColAlgTubes to enable high-yield production of diverse cell types, such as hPSCs-derived endothelial cells^28^, vascular smooth muscle cells^31^, red blood cells^46–48^, platelets^49–52^, beta cells, T cells, and neurons^53–56^. These cells are critical for preclinical and clinical applications and require large-scale production. Expanding ColAlgTubes applications to include other cell populations, such as adult stem cells and immune cells like T cells, will further increase their utility.

## Author Contributions

Y.L. conceived the idea. Y.L., S.W., Y.Y., and X.W. designed the experiments. Y.Y., X.W., and Y.P. conducted the experiments. Y.L., S.W., Y.Y., X.W., Y.P., Y.W., X.L., and A.W. carried out data analysis and manuscript preparation.

## Data Availability

The authors confirm that the data supporting the findings of this study are available within the article.

